# Agent-Based Modeling of Idiopathic Lung Fibrosis and Mechanistic Treatments

**DOI:** 10.64898/2026.03.22.713503

**Authors:** Narshini D. Gunputh, Eirini Kilikian, Claudia A. Miranda, Shayn M. Peirce, Ashlee N. Ford Versypt

## Abstract

Agent-based modeling (ABM) is a computational method for predicting the emergent outcomes of interacting, autonomous individuals in a complex system. Here, ABM is used to simulate interactions between fibroblast and myofibroblast cells during idiopathic pulmonary fibrosis (IPF) in alveolar tissue microenvironments. These microenvironments are derived from histology of a healthy human lung sample and moderate- and severe-IPF lung samples. Fibroblast differentiation, cell migration, and collagen secretion in response to the spatial distribution of the cytokine transforming growth factor-beta are captured in the ABM using NetLogo software. Results are presented from one simulated year without treatment and with mechanisms representing treatment by pirfenidone and pentoxifylline, alone and in combination. A total of 180 *in silico* experiments are run, analyzed, and compared in a high-throughput workflow. The effects of the initial number of fibroblasts and treatment scenarios on various metrics related to collagen accumulation and collagen invasion into alveolar regions are determined. The ABM and the analysis files are shared to facilitate model reuse. By integrating computational modeling of IPF and therapeutics, this research aims to improve understanding of fibrosis progression and assess the efficacy of novel and existing treatments targeting different mechanisms to inform decision-making for IPF treatment.

## 1 Introduction

Idiopathic pulmonary fibrosis (IPF) is a progressive and fatal lung disease that occurs within the alveoli of the lung (Fig. 1). IPF is characterized by excessive extracellular matrix (ECM) deposition in the alveolar interstitial space, leading to destruction of alveoli, increased lung stiffness, and impaired gas exchange. The excessive deposition of ECM is characterized by elevated collagen production by myofibroblasts, which have been activated and differentiated from fibroblasts by growth factors such as TGF-β. In the United States alone, about 100,000 people have been diagnosed with IPF, and the ages of diagnosed people typically range between 50 and 70 years old [1, 2]. The etiology of IPF is not fully understood. The progression of IPF is often due to an imbalanced response to injury that affects tissue remodeling in the lungs [2]. The median survival time of IPF, if left untreated, is 2–3 years after diagnosis [2], and current treatments have limited efficacy [3]. The FDA-approved therapies for IPF, pirfenidone (pirf) and nintedanib, work primarily by slowing disease progression rather than reversing fibrosis. Moreover, the current success rates of both drugs in inhibiting further fibrosis in patients with high fibrosis scores are low [4].

**Fig. 1.**
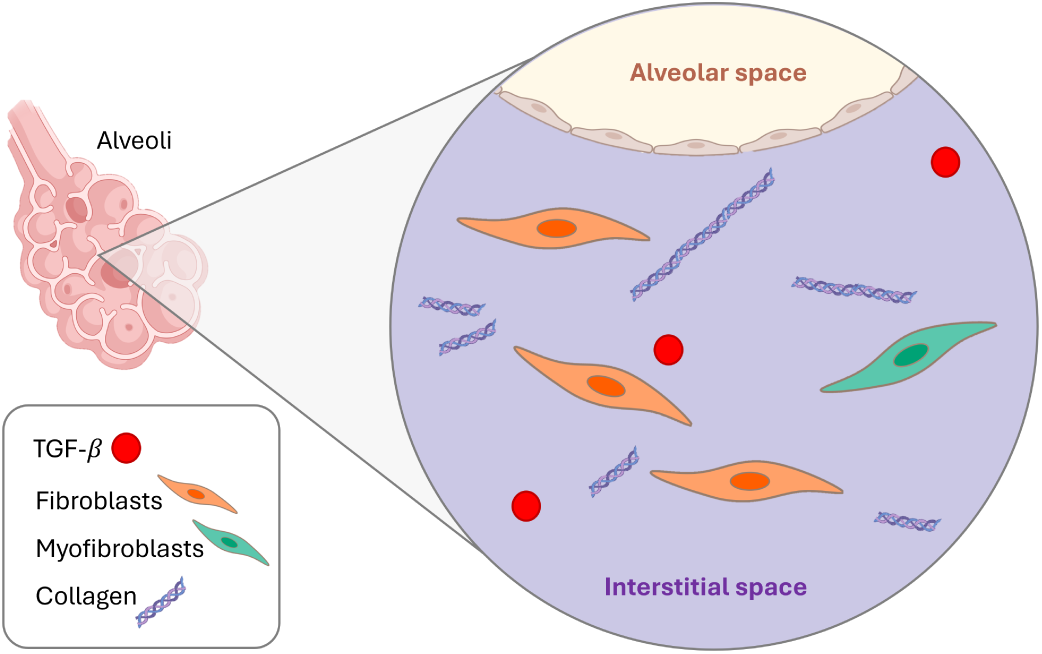
Schematic of the idiopathic pulmonary fibrosis tissue microenvironment within lung alveoli. Created with BioRender.com.

Lung and liver fibrosis share common pathways for fibrosis progression, and researchers have suggested that insights from one organ may inform therapeutic strategies for the other [5–7]. Research on liver fibrosis has identified promising therapeutic agents targeting key fibrotic pathways, including inhibition of transforming growth factor-beta (TGF-β), modulation of macrophages, and ECM remodeling [6, 7]. Pentoxifylline (pentox) has been used in clinical trials for liver fibrosis [8–10]. Pentox treatment effectively reduced pulmonary fibrosis and led to fibroblast cellular senescence in a bleomycin-induced mouse model of IPF [11].

Various mathematical approaches have been used to investigate the complex, multi-scale interactions between cells responsible for fibrosis and their microenvironment during the progression of various forms of lung fibrosis [12]. Mechanistic models that use partial differential equations (PDEs) provide analytical insights into fibroblast proliferation, ECM remodeling, and the secretion of various inflammatory chemokines and can be readily solved using standard numerical methods. However, PDE-based approaches struggle to capture the high spatial and cellular complexity of the lung tissue microenvironment. One such model cleverly addresses spatial organization by homogenizing the tissue across the lung before allowing movement of cells and signaling molecules [13]; however, this approach does not capture the stochastic nature of cellular behaviors or the heterogeneity of the IPF lung epithelium and the available locations for cell movement. To overcome these limitations, agent-based models (ABMs) have been developed to represent cell-cell and cell-tissue interactions, such as proliferation, migration, and differentiation, by incorporating spatial and temporal effects and stochasticity to reflect biological processes more accurately. These models account for individual cellular entities’ interactions to generate realistic and heterogeneous fibrotic patterns in collagen [14–16] and microvascular remodeling [17].

While computational models have enhanced the understanding of the pathophysiology of lung fibrosis, they have been used only to a limited extent to predict drug effects. Our previous work [17] examined the effects of nintedanib on the microvasculature in IPF. A proprietary model for quantitative systems pharmacology applied to IPF has also been described at conferences [18], but no details have been published in the open literature. In this work, we leverage an ABM framework to investigate key cellular and molecular pathways underlying ECM (collagen) deposition in IPF and implement therapeutic interventions via mechanisms of pirf and pentox. By integrating computational modeling of IPF and therapeutic mechanisms, this research aims to improve understanding of fibrosis progression and assess the efficacy of targeting distinct mechanisms to inform decision-making for IPF treatment.

## 2 Methods

To simulate mechanisms of fibrosis and drug effects in the lung alveolar interstitial space (Fig. 1), we developed an ABM focused on the deposition of collagen by fibroblasts and myofibroblasts during the progression of IPF. This *in silico* model is built using NetLogo software [19]. Fibroblasts maintain the structure of connective tissue by secreting collagen, a main component of the ECM. Fibroblasts can differentiate into myofibroblasts [20], which are an activated version of fibroblasts that become highly secretory and contractile and play an important role in wound healing and inflammation [21, 22]. Differentiation is promoted by TGF-β, a cytokine that induces both differentiation and fibrotic collagen production [23–29]. We include fibroblasts, myofibroblasts, TGF-β, and collagen in the ABM. The ABM is fully described in the remainder of this section. The list of agents, adjustable parameters, and outputs for the ABM is provided in Table 1.

**Table 1.**
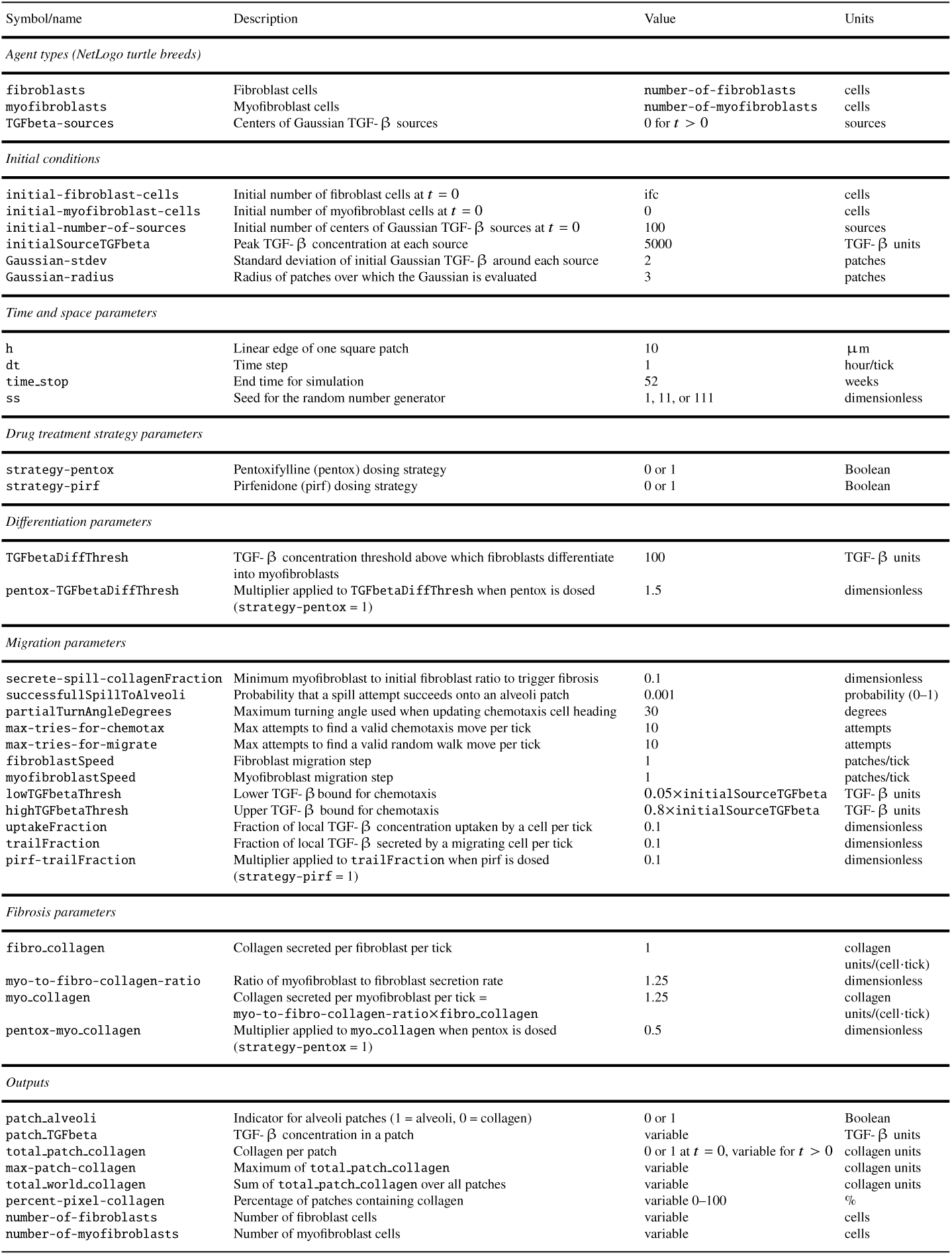
Agents, parameters, and outputs for the NetLogo agent-based model.

### 2.1 Environment

We construct a periodic two-dimensional *in silico* environment or “world” consisting of alveoli (represented by white pixels or “patches” in NetLogo terminology) and ECM primarily composed of collagen (represented by purple patches). The domain is a square of 101 × 101 patches, centered at the origin (i.e., each side ranges from -50 to 50). NetLogo considers patches to be stationary agents with characteristics (i.e., object-oriented attributes), some built-in and some user-defined. We use the built-in characteristics of `pxcor` and `pycor` for each patch’s *𝑥* and *𝑦* coordinates and `pcolor` to indicate whether a patch contains collagen (purple) or alveoli (white). Initial collagen is indicated by light purple (`pcolor` = 117), and patches that switch from alveoli to new collagen deposited during the simulation are indicated by dark purple (`pcolor` = 115). We define a binary Boolean token patch alveoli to indicate whether the patch contains alveoli. Patch characteristics total patch collagen and patch TGFbeta track the dimensionless amounts of collagen and TGF-β on each patch, respectively, and are considered as model outputs of interest. A few additional patch characteristics are defined to hold temporary variables used in calculations to update total patch collagen and patch TGFbeta.

### 2.2 Agents

The simulations consider the behaviors of two types of mobile agents (or “breeds of turtles” as they are called in NetLogo): fibroblasts and myofibroblasts. Myofibroblasts are a more active state of fibroblasts, meaning they are more aggressive in their interactions with TGF-β and collagen secretion. Myofibroblasts have the same behaviors as fibroblasts, except for differentiation, which is an exclusive feature of fibroblasts. Each agent type has different values for user-defined properties (Table 1). Each agent (i.e., turtle) has a directional heading, *𝑥* and *𝑦* coordinates, a shape icon, a breed, and a size. We use custom shape icons to visualize fibroblasts and myofibroblasts. Each agent also has custom characteristics to hold temporary variables for migration rules and agent interactions with TGF-β.

A third type of agent TGFbeta-sources is used to initialize hotspots of TGF-β representing diffuse injury of unknown origin to exacerbate IPF. Immediately after placement and update of the corresponding patch variable patch TGFbeta, these sources are removed and no longer considered as agents. The placement of initial TGFbeta-sources is described further in Sect. 2.3.

### 2.3 Initialization

The virtual environment for the NetLogo world is built from human lung tissue sections stained with hematoxylin and eosin (H&E). These tissue samples were collected, studied, and shared by Franzén et al. [30]. The H&E-stained tissue sections are marked with fiducial frames for alignment to Visium spatial transcriptomics; thus, the images are consistently 8 mm × 8 mm. To focus on the interior of the tissue and avoid edges of the tissue sections, we select the center one-eighth of the tissue image regions (1000 µm × 1000 µm for the domain). We aim to capture 20 µm-scale features, so we specify that each NetLogo patch is 10 µm (5 pixels) on a side. Thus, each pixel in NetLogo is 2 µm. In a square domain of 1000 µm on a side, the H&E stain images have 1500 pixels on a side, giving a scaling of 1.5 image pixels per 1 µm. We use the software ImageJ [31] with the following script to automate the image processing to set the scale, select the centered region of interest, crop the domain, and binarize the resulting image with a black and white mask:

`run(“Set Scale…”,“distance=1.5 known=1 unit=unit”);`

`run(“Specify…”,“width=1000 height=1000 x=3500 y=3500 scaled”);`

`run(“Crop”);`

`setOption(“BlackBackground”,false);`

`run(“Convert to Mask”);`

Then we open the output image from the ImageJ batch script in our NetLogo code. We run the following command to convert the black pixels to our desired shade of purple: `patches> if pcolor != 9.9 [set pcolor 117`]. We save the corresponding NetLogo world as a .csv file so we can easily load the domain for each tissue section. This process is applied to all 25 lung tissue samples from [30], and the NetLogo worlds are available in our GitHub repository [32]. Here, we use the high-resolution images obtained from lung resections from one healthy subject (sample HC 3.B0.2/V10T03-280-A1, which we refer to as case A) and one patient with IPF, from which tissue blocks of different severity of fibrotic injury were studied in [30]. From the IPF patient samples, we select a moderate fibrosis-graded section (sample IPF 1.B2.1/V19S23-092-C1, which we refer to as case C) and a severe fibrosis tissue section (sample IPF 1.B3.2/V10T03-279-D1, which we refer to as case D). Mild fibrosis-graded sections corresponding to case B exist in the data set, but we did not select this case to process further here.

Where a patch is collagen (`patch alveoli` = 0), `total patch collagen` is initialized to 1. Where a patch is alveoli (`patch alveoli` = 1), `total patch collagen` is initialized to 1 (i.e., no collagen on the alveoli patches). The value of patch TGFbeta on alveoli patches is 0.

Within the representative tissue samples, initial agent placement is confined to the collagen patches. The initial number of fibroblasts (ifc) is defined by the variable `initial-fibroblast-cells`. At *𝑡* = 0, the fibroblasts are placed in the center of randomly selected collagen patches. We use ifc in the range of 20 to 100 cells in the domain based on the value of about 60 fibroblasts/mm2 at 7 days from an *ex vivo* culture model for human lung fibrogenesis in IPF [28]. No myofibroblast cells are initially present in the simulation; their only source is through differentiation of fibroblasts during the simulation. Stationary sources of TGF-β (initial-number-of-sources is the user-defined number of sources) are randomly distributed among the collagen patches. The initial TGF-β deposits represent spontaneous activation of this molecule in the ECM due to epithelial cell death [33, 34] or other tissue damage causes, such as low pH or proteolysis [35]. These TGF-β deposits are placed as TGFbeta-source agents on randomly selected collagen patches, independent of the selection of fibroblast initial locations. The coordinates (*𝜇_𝑥_, 𝜇_𝑦_*) of each TGFbeta-source agent are used as the center of a scaled 2D Gaussian function (radially symmetric bell curve):

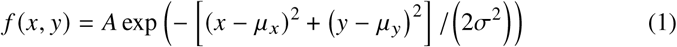

where *𝑓* (*𝑥, 𝑦*) is the amount added to patch TGFbeta in the patch at (*𝑥, 𝑦*) within a radius of Gaussian-radius number of patches from the center with standard deviation of *𝜎* = Gaussian-stdev and amplitude *𝐴* = initialSourceTGFbeta. Any alveoli patches (patch alveoli = 1) within a radius of Gaussian-radius number of patches from the TGFbeta-source agent are ignored and kept at patch TGFbeta = 0. Multiple TGFbeta-source agents may be placed near each other, such that patch TGFbeta values reflect the summed contributions of all nearby sources. At the end of the initialization step, the TGFbeta-source agents are removed from the simulation and not used further; instead, patch TGFbeta is tracked at each patch. The intention behind this choice of functional form for TGF-β initialization is to represent sources diffusing outward over a long time scale, avoiding the need to simulate fast molecular diffusion from recurrent point sources. The values of the parameters for initialization are provided in Table 1.

### 2.4 Rules

For each tick or time step (1 hour of simulated time), the following ABM rules (Fig. 2) are executed, the patch and agent variables are incremented, and each dynamic output metric is updated and recorded. The simulation terminates after *𝑡* = 52 weeks = 8736 ticks. The rules are organized into three sets: differentiation, migration, and fibrosis, which are detailed in turn.

**Fig. 2.**
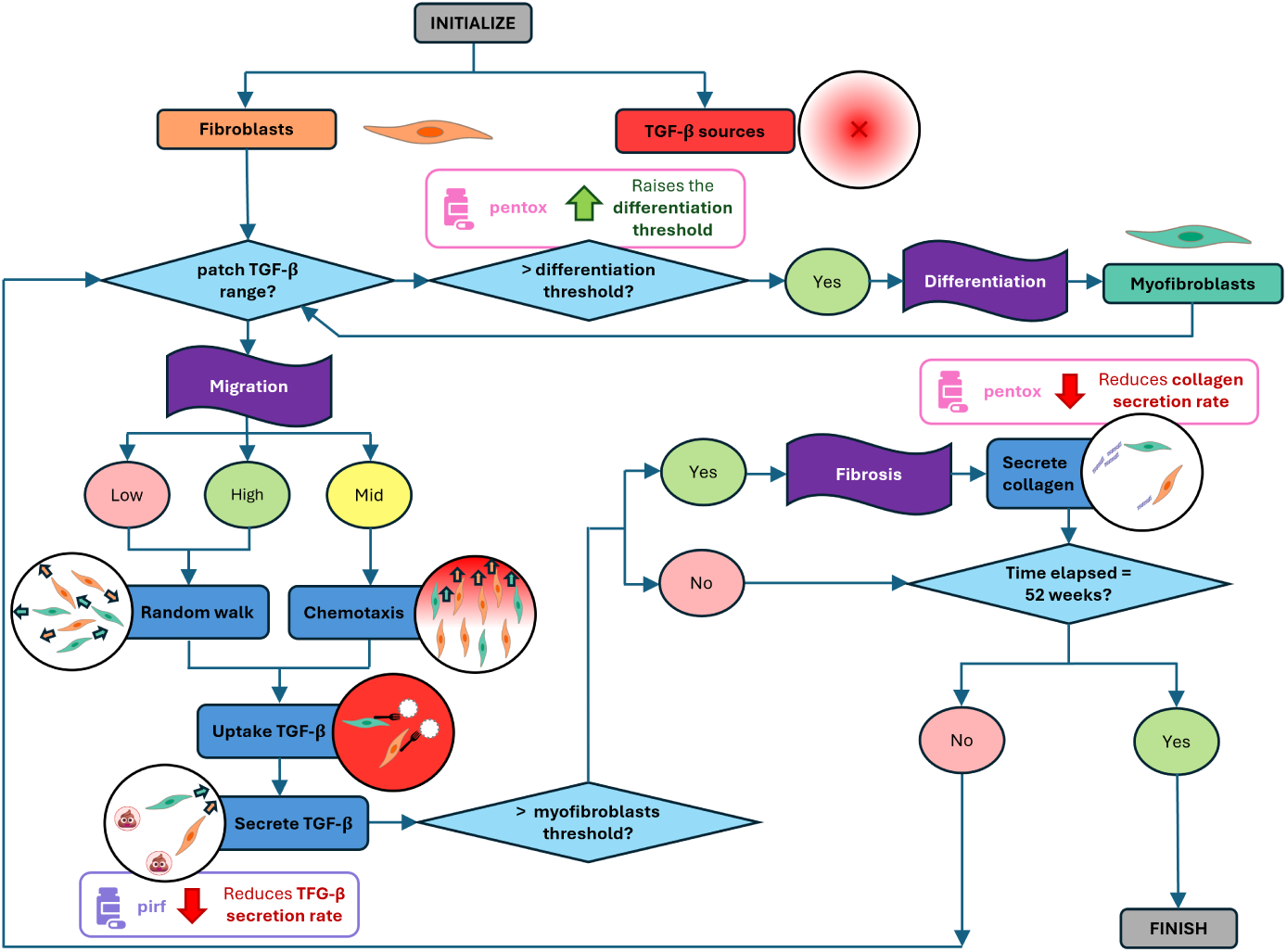
Decision flow chart for the agent-based model for fibroblasts, TGF-β sources, and myofibroblasts interacting in a domain with collagen and alveoli patches via three sets of rules for differentiation, migration, and fibrosis and effects of drug treatment strategies for pentox and pirf.

#### 2.4.1 Differentiation

Each fibroblast checks patch TGFbeta of the patch where it is located to test whether the local patch TGFbeta exceeds the differentiation threshold TGFbetaDiffThresh. If so, the fibroblast switches to the myofibroblast type, and the counts of both cell types are updated. Some parameters depend on cell type to reflect the more aggresive secretory role of myofibroblasts. Also, note that the fibrosis rule is only triggered when the number of myofibroblasts exceeds a specified fraction (secrete-spill-collagenFraction of the initial number of fibroblasts). This is intended to capture that fibrosis happens after fibroblasts are activated by a stimulus (here, TGF-β). Thus, multiple successful calls to the differentiation rule are required before the fibrosis rule is called.

#### 2.4.2 Migration

Each fibroblast and myofibroblast cell enters the decision flow chart for the migration rules once during each time step. The model considers two modes of migration: random walk and chemotaxis. If the local concentration of TGF-β (patch TGFbeta) is low or high, cells move in a random walk fashion (random new angle of movement) on collagen patches at a speed of fibroblastSpeed or myofibroblastSpeed. For patch TGFbeta values between lowTGFbetaThresh and highTGFbetaThresh, cells chemotax by moving toward the adjacent neighbor patch with the highest patch TGFbeta, provided that value exceeds the current patch’s patch TGFbeta. The heading for cell movement is set to move up the local TGF-β gradient, then adjusted to a random direction within ±partialTurnAngleDegrees of that heading.

The bounds for chemotaxis are selected as multipliers of initialSourceTGFbeta. Chemotaxis also proceeds at the same speeds as for a random walk. If migration attempts move a cell onto an alveoli patch, the movement is rejected, and the migration is attempted again in the same time step. This is repeated up to a maximum of max-tries-for-migrate for random walk and max-tries-for-chemotax for chemotaxis, respectively. If the last attempt fails to yield a collagen patch as the destination, the cell does not move and exits the migration rule. The maximum number of attempts is implemented to prevent infinite loops that can result if cells are placed on isolated collagen patches surrounded by alveoli.

The model simulates TGF-β metabolism [36] and cellular interaction via two steps: uptake and secretion. Cells remove uptakeFraction×patch TGFbeta from patch TGFbeta of the patches where they start before migration. After successful migration, cells secrete TGF-β on their destination patches by adding trailFraction×patch TGFbeta to patch TGFbeta.

#### 2.4.3 Fibrosis

The fibrosis rule is called if the number of myofibroblasts is greater than or equal to secrete-spill-collagenFraction×ifc. Once this threshold of myofibroblasts is surpassed, all cells secrete collagen at rates of fibro collagen and myo collagen for fibroblasts and myofibroblasts, respectively. The patch variable total patch collagen is updated by the contributions from each cell on the patch and from cells on adjacent neighboring patches. This spillage to neighboring patches is meant to capture limited local collagen diffusion in the initial soluble form and cross-linking as fibrotic scarring develops. The spill to a neighboring patch is always successful if that patch already contains collagen (i.e., patch alveoli = 0). Cells secrete collagen into adjacent alveoli patches at a success rate of successfullSpillToAlveoli implemented stochastically by comparison to a random number drawn between 0 and 1. If unsuccessful, the patch remains alveoli with patch alveoli = 1 and total patch collagen = 0. If successful, then patch alveoli is switched to 0, and total patch collagen increments by the amount secreted from the adjacent cell. For visualization purposes, the patch color is set to a dark shade of purple, denoting “new” invasive collagen secretion. The value of successfullSpillToAlveoli = 0.001 is used to model a slow invasion rate that takes years to progress from inflammatory injury in a healthy lung to a moderate stage of fibrosis.

### 2.5 Drug treatment strategies

We considered four possible strategies, indicated by the combinations of Boolean parameters strategy-pirf and strategy-pentox, where 0 means no drug is administered, and 1 means the drug is administered at time *𝑡* = 0 and sustained at a constant level.

In animal models of lung fibrosis and human lung cell cultures [11, 37], pentox was shown to reduce fibrosis by inhibiting fibroblast proliferation and reducing collagen production by fibroblasts. Thus, our simulated pentox treatment directly acts through the differentiation and fibrosis rules. In lung cell culture studies [38–40], pirf was also shown to inhibit fibroblast activation and reduce collagen production [6]. Additionally, pirf was shown to prevent fibroblast migration in culture [40]. As it is FDA-approved for patient use in IPF, pirf efficacy in humans has also been established [41, 42]. The mechanism is not fully elucidated for pirf in humans, but the effects of pirf are mediated by suppression of TGF-β [29, 43, 44]. Thus, we model pirf treatment as impacting TGF-β and, indirectly, fibrosis via downstream effects through the differentiation and fibrosis rules. The scenario of having both pirf and pentox treatment considers the combination of indirect action on fibrosis through migration-rule effects on TGF-β secretion and direct actions via differentiation and fibrosis rules.

Our choice on how to apply treatment strategies *in silico* comes from the available literature on the possible molecular pathways these drugs may affect. However, the data remain inconclusive about their specific overall mechanisms. Thus, one should consider our simulated “pentox” treatment as modulating the fibrosis and differentiation rules and our simulated “pirf” treatment as directly perturbing an aspect of the migration rule related to TGF-β secretion (Fig. 2 and Table 1). Specifically, pentox increases TGFbetaDiffThresh for differentiation by the multiplier pentox-TGFbetaDiffThresh and reduces the collagen secretion rate from myofibroblasts through pentox-myo collagen multiplied by myo collagen. Pirf reduces trailFraction for TGF-β secretion after successful migration by the multiplier pirf-trailFraction.

### 2.6 *In silico* experiments and output metrics

We implement a streamlined series of *in silico* experiments using the BehaviorSpace functionality in NetLogo and the human lung tissue samples (cases A, C, and D). BehaviorSpace enables high-throughput automation of *in silico* experiments across combinations of multiple input settings, allowing for a thorough parameter sweep. For each tissue sample case, the NetLogo simulation is run for 52 weeks for the following set of inputs across each combination of initial number of fibroblast cells, pentox treatment strategy, pirf treatment strategy, and replicates starting at different random seeds:

- initial-fibroblast-cells: ifc = {20, 40, 60, 80, 100}
- strategy-pentox: {0, 1}
- strategy-pirf: {0, 1}
- random seed: ss = {1, 11, 111}

The output files for each of the 180 runs include spreadsheets with agent and patch variables at the beginning and end of the run and images of the NetLogo world showing all fibroblasts and myofibroblasts, the alveoli patches, and collagen patches at the beginning and end of the run. For each tissue sample case (a BehaviorSpace set of *in silico* experiments), a single file called “spreadsheet.csv” is generated that records the following time-dependent scalar reporter metrics:

- total world collagen: referred to as total collagen and determined by summing total patch collagen over all patches
- percent-pixel-collagen: percentage of patches in the domain containing collagen, determined by taking the ratio of patches with patch alveoli = 0 to the total number of patches, and referred to as percent collagen
- number-of-fibroblasts: number of fibroblast cells in the domain
- number-of-myofibroblasts: number of myofibroblast cells in the domain
- max-patch-collagen: maximum intensity of collagen in the domain, determined by taking the maximum of total patch collagen, and referred to as max collagen

## 3 Results and Discussion

### 3.1 NetLogo simulations for lung tissue cases and treatment conditions

For each 52-week simulation, we collected the NetLogo world’s initial and final spatial configurations. The first row of Fig. 3 shows these configurations for replicate 1 of case A healthy lung tissue for the control scenario without treatment. For each patch in the domain, we have the magnitudes of the amounts of collagen and TGF-β from the patch characteristics total patch collagen and patch TGFbeta, respectively. For each cell, we have the coordinates of its center at the initial and final times. We used MATLAB to render contour maps of total patch collagen (Fig. 3 second row) and TGF-β normalized by the peak value for a single source (i.e., patch TGFbeta/initialSourceTGFbeta for Fig. 3 third row).

**Fig. 3.**
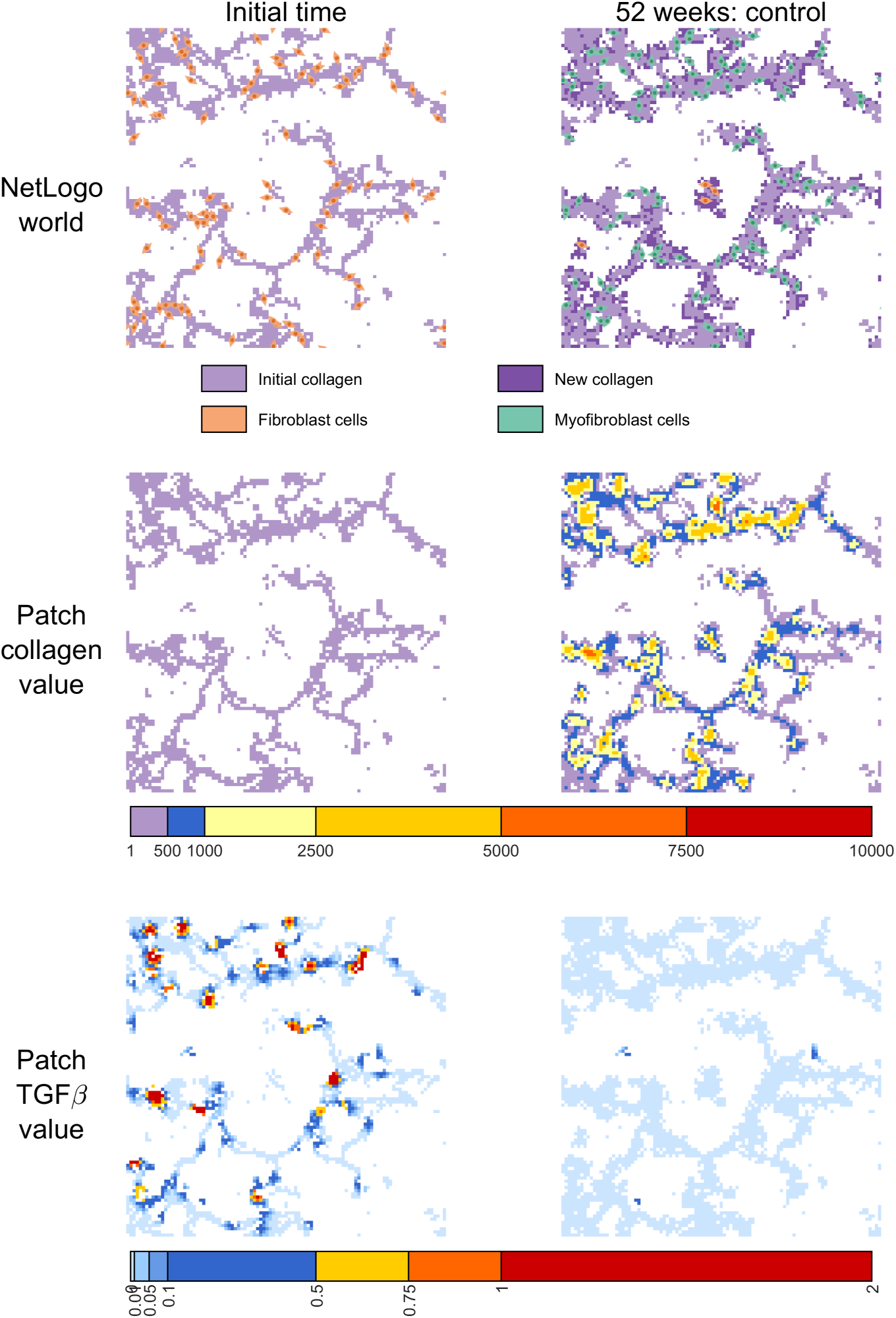
Spatial data for *in silico* experiment replicate 1 (random seed = 1) for case A healthy lung tissue for 52 weeks without treatment. (First column): spatial data at the initial time. (Second column): spatial data at 52 weeks of simulated time. (First row): NetLogo world spatial configurations. Light purple patches show initial collagen, dark purple patches indicate new collagen deposited during the simulation, fibroblast cells are orange, and myofibroblast cells are green. (Second row): contour maps of total patch collagen (collagen units). (Third row): contour maps of patch TGFbeta/initialSourceTGFbeta (normalized TGF-β units).

Our simulations considered four scenarios of treatment for 52 weeks: control, pirf, pentox, and pentox & pirf. The visualization in Fig. 3 is extended to show the treatment scenarios in adjacent columns in Fig. 4a. This view is our standard for visually comparing spatial data for *in silico* experiments. We repeated the simulations shown in Fig. 4a for two more replicates with varying initial configurations of locations for fibroblast cells and TGF-β sources. Note that all treatment scenarios for a given replicate have the same initial configurations. Results for replicate 2 with random seed = 11 and replicate 3 with random seed = 111 are shown in Fig. 4b,c, respectively. We repeated the process for case C moderate fibrosis and case D severe fibrosis (Figs. 5 and 6). Fig. 7 shows the NetLogo worlds for treatments across replicates to facilitate comparisons for each case. Note that we do not consider mild fibrosis (case B) in our analysis presented here.

**Fig. 4.**
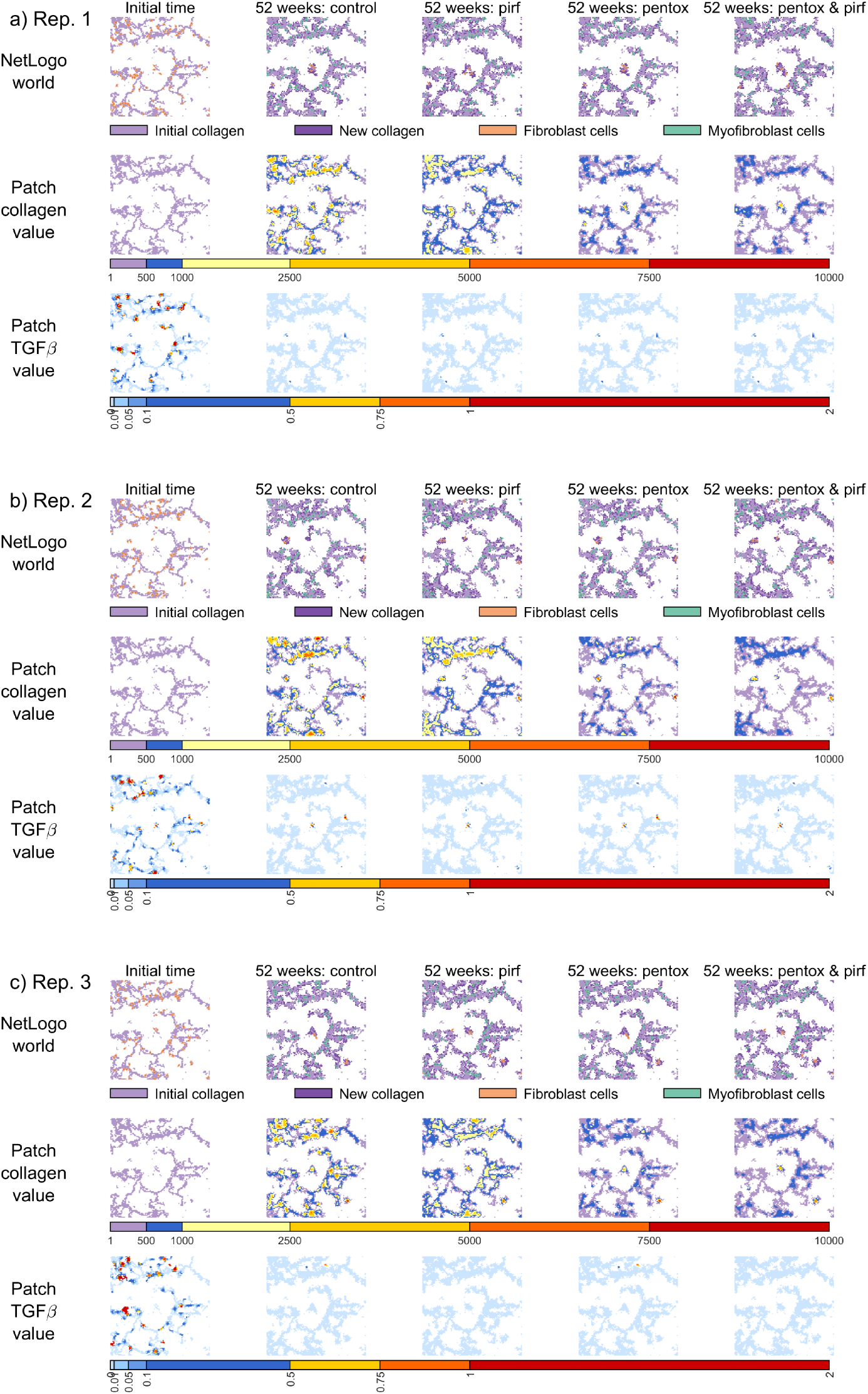
Spatial data for *in silico* experiments for case A healthy lung tissue for 52 weeks of simulated time with initial fibroblast cell and TGF-β source locations varying by replicate (Rep.). (First column): spatial data at the initial time. (Second column): spatial data at 52 weeks without treatment. (Third column): spatial data at 52 weeks with pirf treatment. (Fourth column): spatial data at 52 weeks with pentox treatment. (Fifth column): spatial data at 52 weeks with pentox and pirf treatment. (First row): NetLogo world spatial configurations. Light purple patches show initial collagen, dark purple patches indicate collagen new deposited during the simulation, fibroblast cells are orange, and myofibroblast cells are green. (Second row): contour maps of total patch collagen (collagen units). (Third row): contour maps of patch TGFbeta/initialSourceTGFbeta (normalized TGF-β units). a) Rep. 1 (random seed = 1). b) Rep. 2 (random seed = 11). c) Rep. 3 (random seed = 111).

**Fig. 5.**
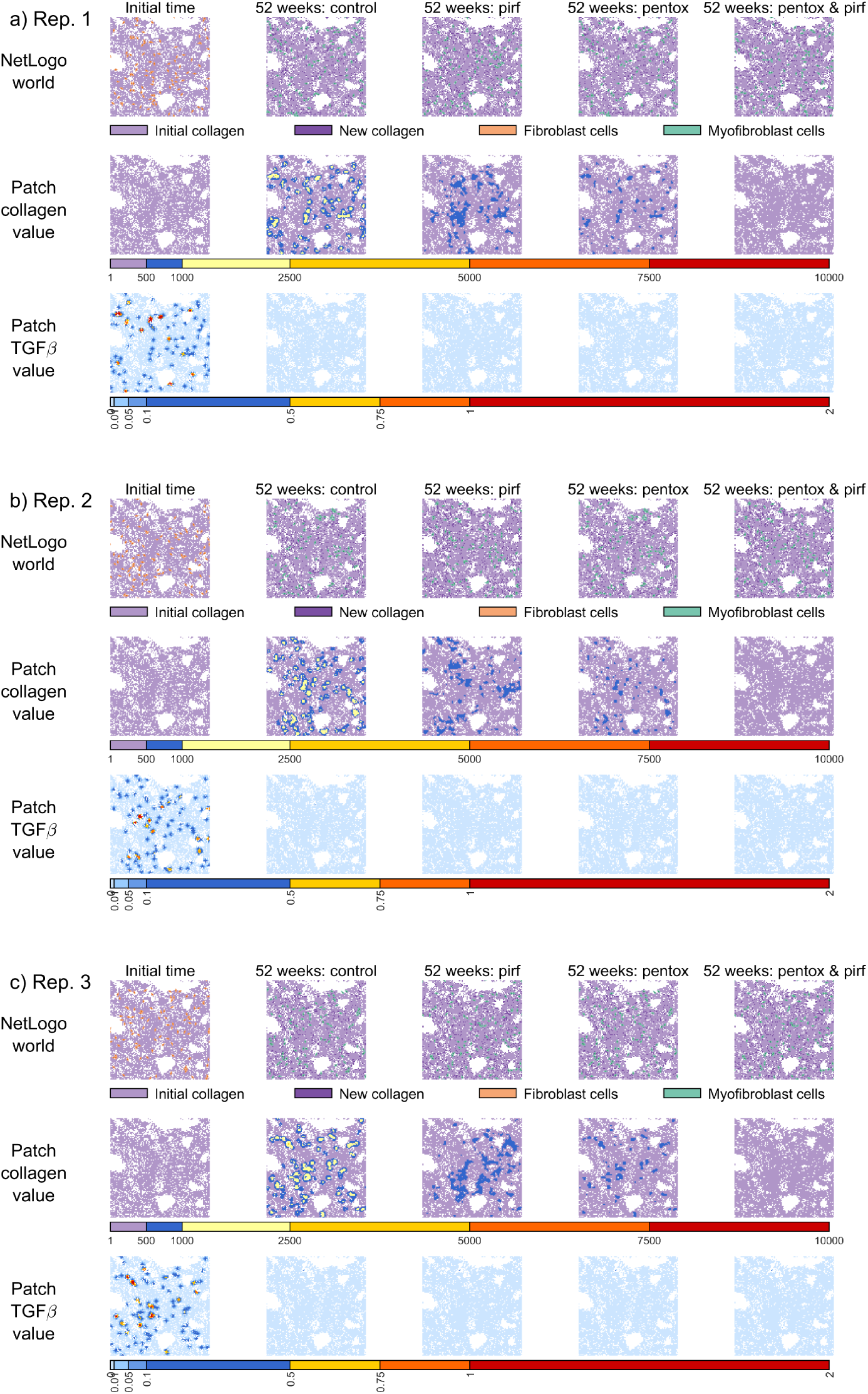
Spatial data for *in silico* experiments for case C moderate fibrosis lung tissue for 52 weeks of simulated time with initial fibroblast cell and TGF-β source locations varying by replicate (Rep.). (First column): spatial data at the initial time. (Second column): spatial data at 52 weeks without treatment. (Third column): spatial data at 52 weeks with pirf treatment. (Fourth column): spatial data at 52 weeks with pentox treatment. (Fifth column): spatial data at 52 weeks with pentox and pirf treatment. (First row): NetLogo world spatial configurations. Light purple patches show initial collagen, dark purple patches indicate new collagen deposited during the simulation, fibroblast cells are orange, and myofibroblast cells are green. (Second row): contour maps of total patch collagen (collagen units). (Third row): contour maps of patch TGFbeta/initialSourceTGFbeta (normalized TGF-β units). a) Rep. 1 (random seed = 1). b) Rep. 2 (random seed = 11). c) Rep. 3 (random seed = 111).

**Fig. 6.**
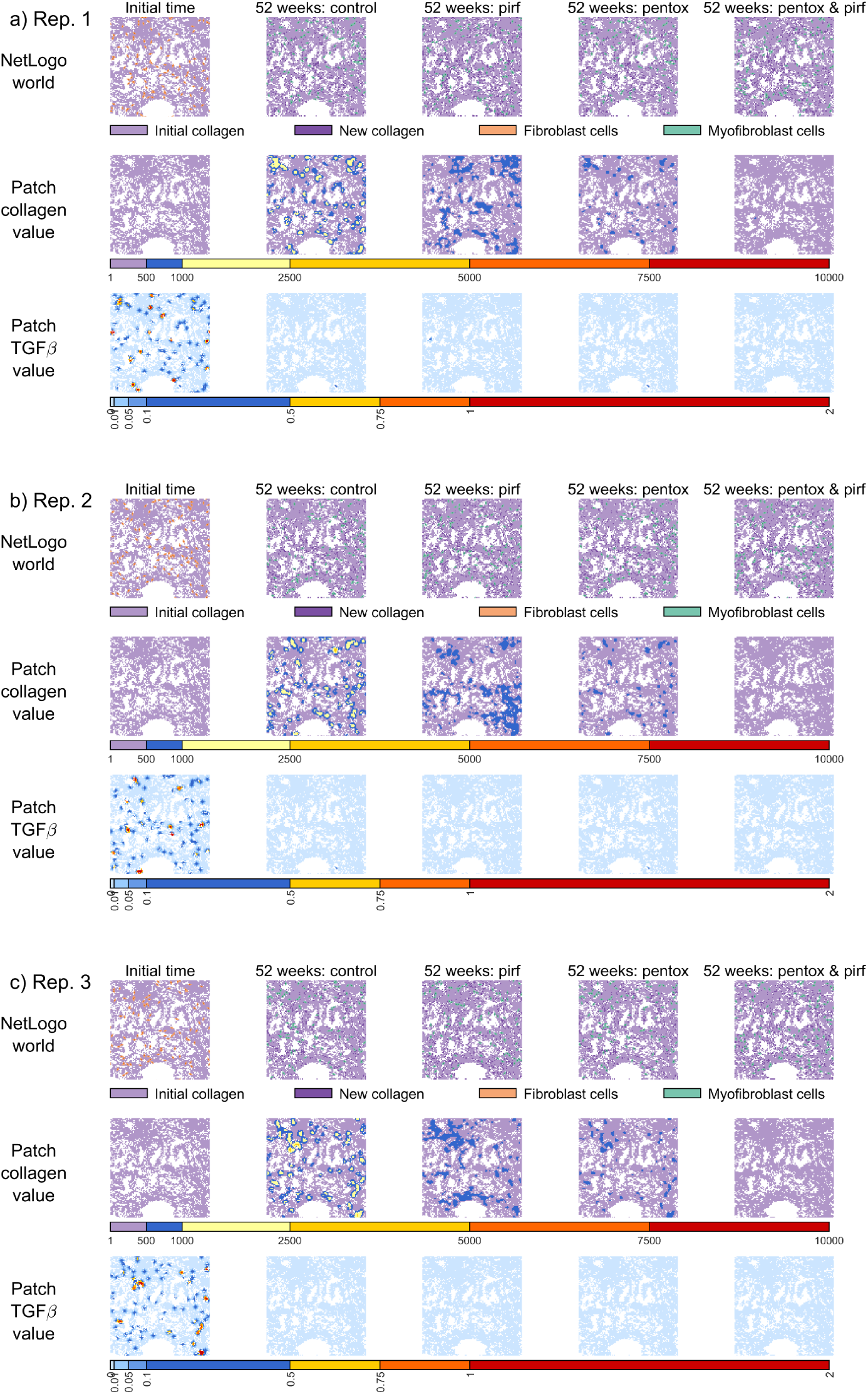
Spatial data for *in silico* experiments for case D severe fibrosis lung tissue for 52 weeks of simulated time with initial fibroblast cell and TGF-β source locations varying by replicate (Rep.). (First column): spatial data at the initial time. (Second column): spatial data at 52 weeks without treatment. (Third column): spatial data at 52 weeks with pirf treatment. (Fourth column): spatial data at 52 weeks with pentox treatment. (Fifth column): spatial data at 52 weeks with pentox and pirf treatment. (First row): NetLogo world spatial configurations. Light purple patches show initial collagen, dark purple patches indicate new collagen deposited during the simulation, fibroblast cells are orange, and myofibroblast cells are green. (Second row): contour maps of total patch collagen (collagen units). (Third row): contour maps of patch TGFbeta/initialSourceTGFbeta (normalized TGF-β units). a) Rep. 1 (random seed = 1). b) Rep. 2 (random seed = 11). c) Rep. 3 (random seed = 111).

**Fig. 7.**
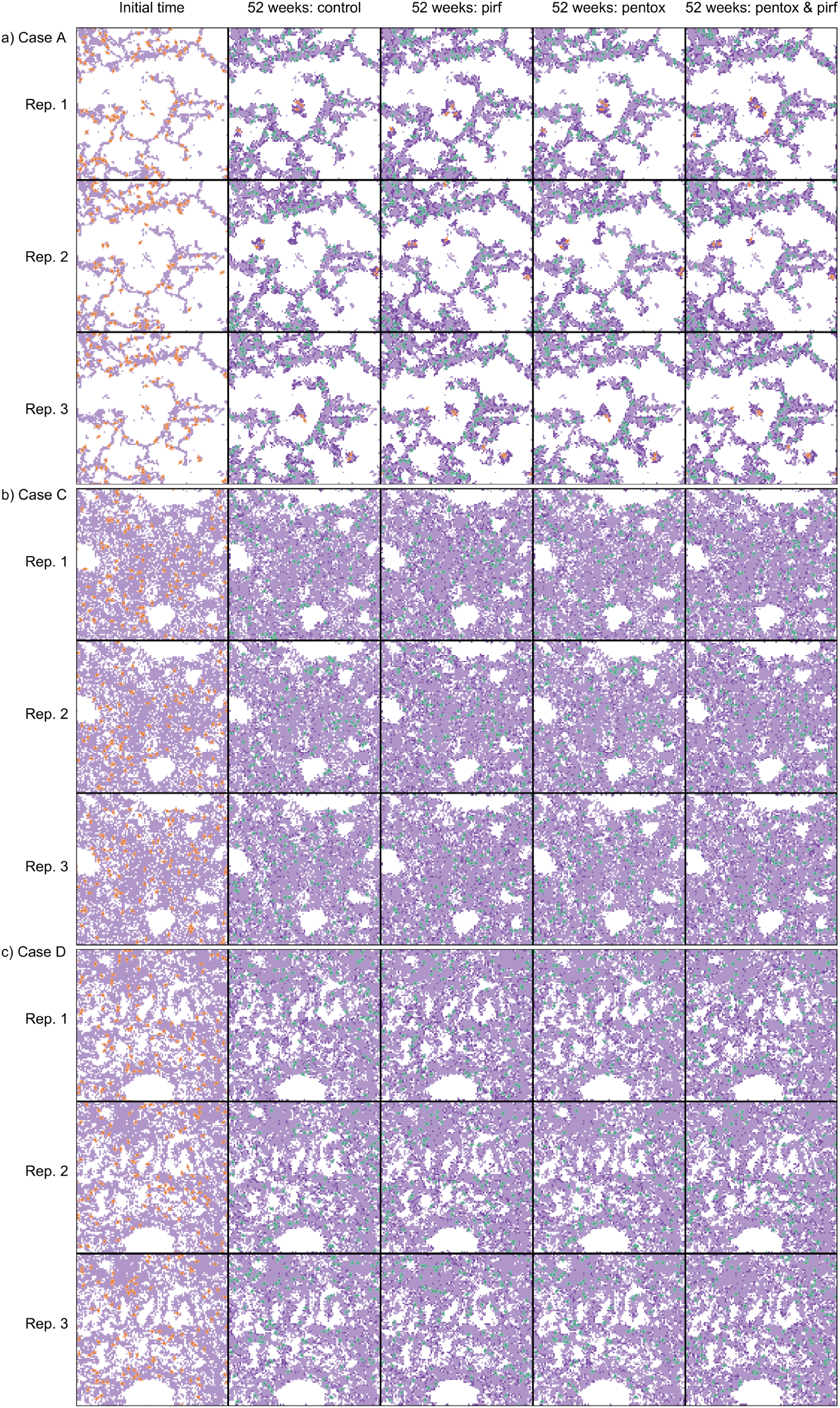
NetLogo world spatial configurations for *in silico* experiments for a) case A healthy, b) case C moderate fibrosis, and c) case D severe fibrosis lung tissue for 52 weeks of simulated time with initial fibroblast cell and TGF-β source locations varying by replicate (Rep.). Light purple patches show initial collagen, dark purple patches indicate new collagen deposited during the simulation, fibroblast cells are orange, and myofibroblast cells are green. (First column): spatial data at the initial time. (Second column): spatial data at 52 weeks without treatment. (Third column): spatial data at 52 weeks with pirf treatment. (Fourth column): spatial data at 52 weeks with pentox treatment. (Fifth column): spatial data at 52 weeks with pentox and pirf treatment. (First row): Rep. 1 (random seed = 1). (Second row): Rep. 2 (random seed = 11). (Third row): Rep. 3 (random seed = 111).

### 3.2 Reporter metrics to compare simulation results

All of the simulation results shown in Figs. 3–7 in Sect. 3.1 had 100 initial fibroblast cells (i.e., ifc = 100). We simulated the treatment conditions for all three tissue sample cases, each with three replicates and ifc = {20, 40, 60, 80, 100}. To allow for more than visual inspection of Figs. 3–7 or their counterparts across the range of ifc (a total dataset of 180 simulations), we calculated and tracked the five summary reporter metrics defined in Sect. 2.6 at every time step for each NetLogo simulation. Throughout the remainder of this section, colors are used to denote values of ifc. Line styles are used to denote treatment conditions: control, pirf, pentox, and pentox & pirf. Fig. 8 displays the five metrics over time for all simulations in this study. Each of these metrics is analyzed and discussed in turn.

**Fig. 8.**
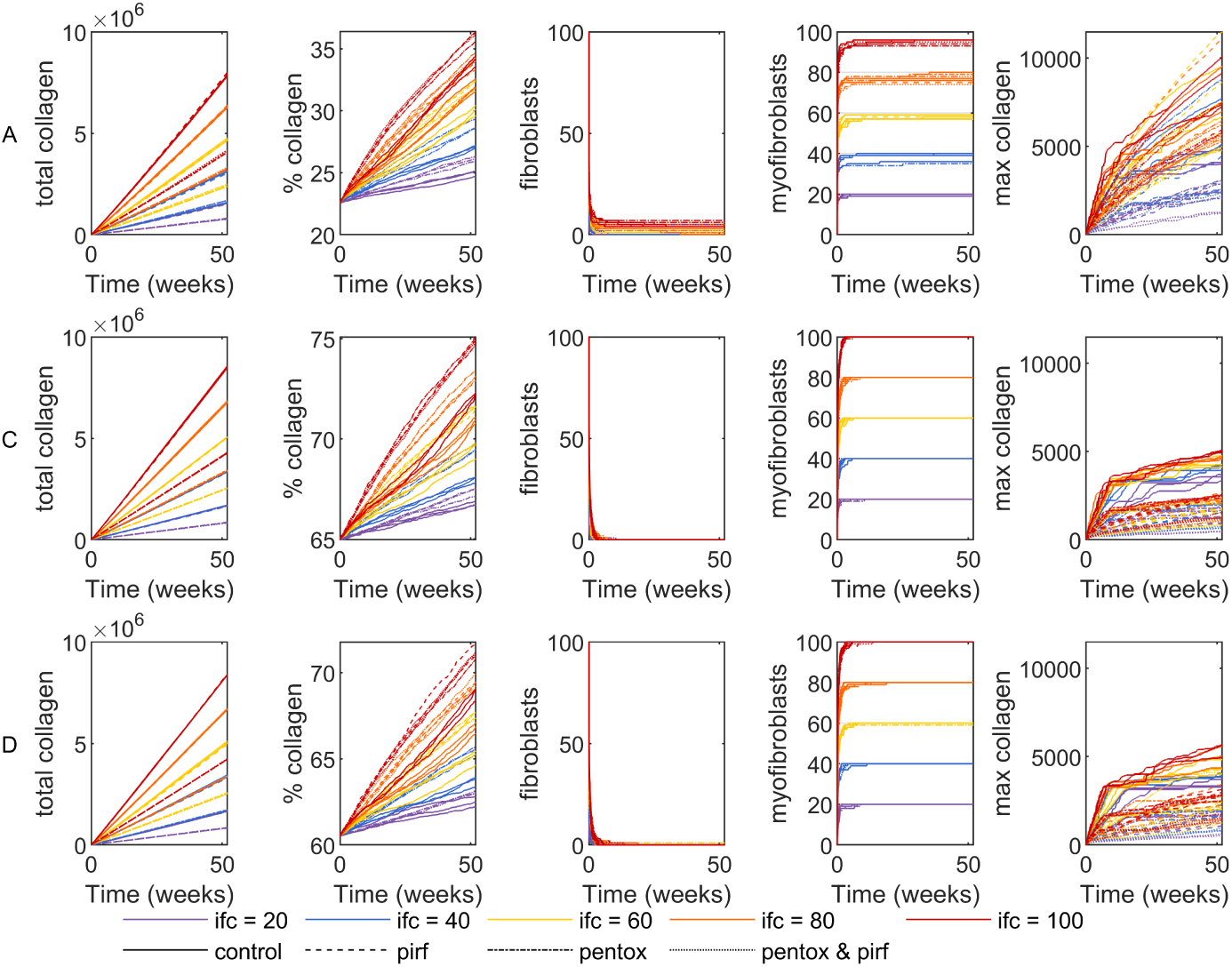
Dynamic metrics for 180 *in silico* experiments automated through BehaviorSpace in NetLogo for (first row) case A healthy, (second row) case C moderate fibrosis, and (third row) case D severe fibrosis lung tissue for 52 weeks of simulated time. Colors are used to denote values of initial-fibroblast-cells = ifc = {20, 40, 60, 80, 100}. Line styles are used to denote treatment conditions: control, pirf, pentox, and pentox & pirf. (First column): total collagen. (Second column): percent collagen. Note the differences in scales between each row. (Third column): number of fibroblast cells. (Fourth column): number of myofibroblast cells. (Fifth column): max collagen.

#### 3.2.1 Numbers of cells

Because the metrics of the numbers of fibroblasts and myofibroblasts demonstrate dependence primarily on ifc, we only briefly discuss these metrics from the curves in Fig. 8. For all simulations, the number of fibroblasts quickly approaches 0. For some simulations, the final number of fibroblasts stabilizes at a small, nonzero number. This is due to isolated instances where an initial fibroblast is placed on a collagen “island” (i.e., a region that is disconnected from the other collagen regions) that is not initialized with any nearby TGF-β that can stimulate the fibroblast to differentiate into a myofibroblast. The island is sufficiently remote that collagen deposition during the simulation never connects it to other collagen regions in the domain. These situations occur for all three replicates of case A as a small number of fibroblasts are placed in such a manner (in Figs. 3 and 4 compare the locations of the remaining fibroblast cells after 52 weeks to the corresponding initial patch TGFbeta). This occurs more frequently when ifc is higher (i.e., Fig. 8 where fibroblasts for case A have larger nonzero final values for red curves compared to orange curves, compared to yellow curves, and so on). Because the collagen regions in cases C and D are more interconnected, no instances of isolated fibroblasts persist throughout the simulations for ifc = 100 (in Fig. 7 notice the lack of orange fibroblasts in any 52-week treatment condition or replicate for case C or case D compared to those for case A). The number of myofibroblasts is simply the difference between ifc and the number of fibroblast cells. Thus, the myofibroblast dynamics (Fig. 8) display the consequences of the isolated fibroblasts clearly. For case A, the difference between ifc (the *𝑦*-axis ticks) and the final myofibroblasts increases with ifc. For case C, all simulations yield the final number of myofibroblasts equal to ifc. For case D, all but two of 60 simulations yield the final number of myofibroblasts equal to ifc as in case C. The two simulations with 1 isolated fibroblast at 52 weeks for case D are replicate 2 of the pirf and pentox & pirf treatments.

#### 3.2.2 Total collagen

The total collagen metric captures collagen deposition but not spatial heterogeneity. The total patch collagen values of some scenarios are quite heterogeneous at 52 weeks, particularly those without drug treatment (Figs. 3–6). The first column of Fig. 8 shows the total collagen dynamics for cases A, C, and D. For alternative comparisons that spread the 60 simulations for each case into five panels organized by ifc and by the treatment scenarios, Fig. 9 shows the total collagen dynamics. The gray lines in Fig. 9 indicate reference trajectories of constant slope of 10×ifc/tick and 5×ifc/tick. These were determined by inspection and were not expected in advance. The total collagen dynamics appear linear in all simulations. However, slight nonlinearities exist due to stochastic fluctuations and subtle slope shifts as the cell population transitions from fibroblasts to myofibroblasts secreting at different rates (Fig. 9). Collagen deposition happens at every time step at rates specific for each cell type and treatment condition. Cases C and D start with more connected collagen regions, thus they are less likely than case A to result in isolated fibroblasts or isolated TGF-β sources upon random initialization, and they more consistently approach the total collagen increase rate of 10×ifc/tick for the control scenario.

**Fig. 9.**
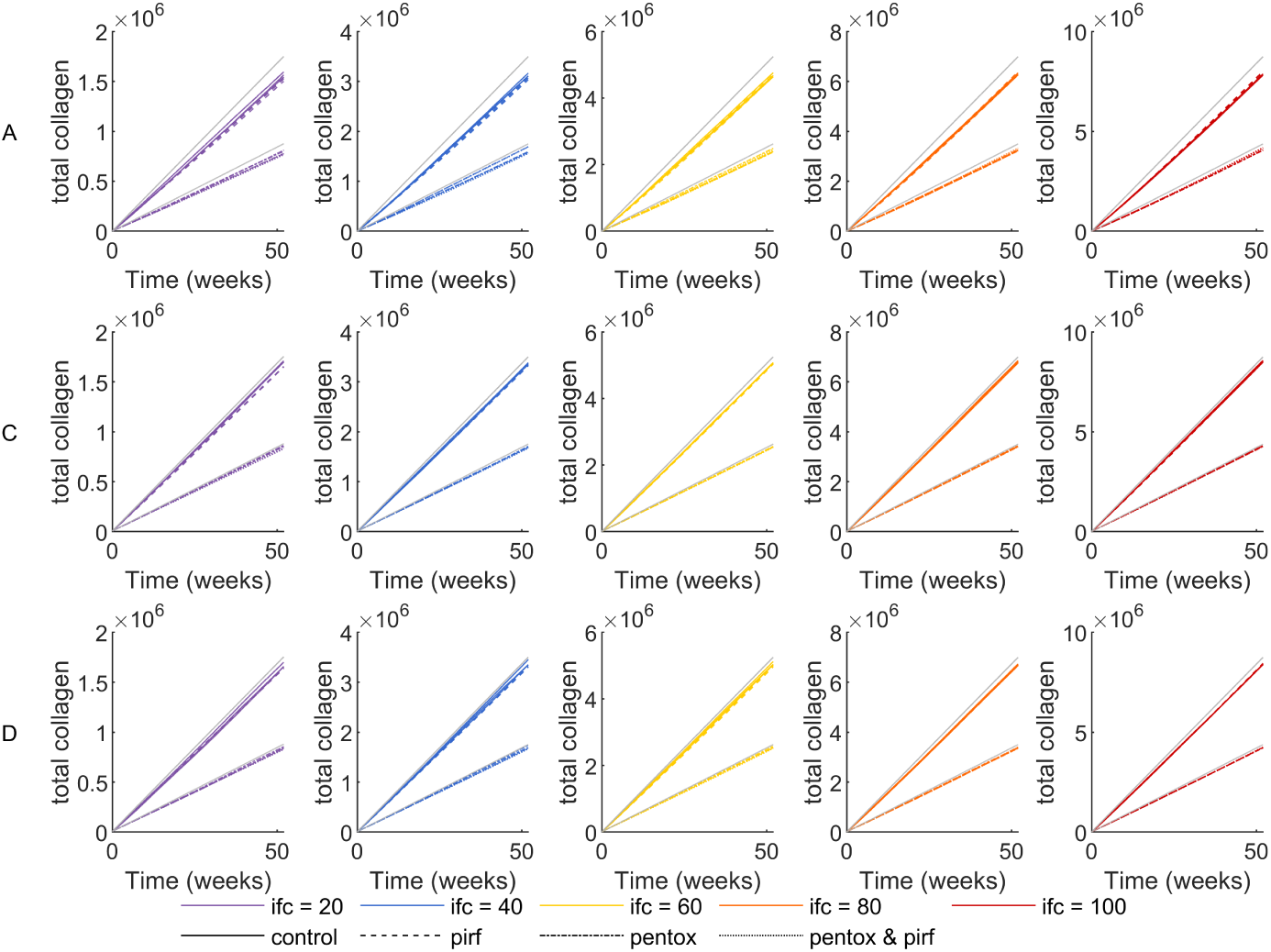
Dynamic total collagen for 180 *in silico* experiments automated through BehaviorSpace in NetLogo for (first row) case A healthy, (second row) case C moderate fibrosis, and (third row) case D severe fibrosis lung tissue for 52 weeks of simulated time. Colors are used to denote values of initial-fibroblast-cells = ifc = {20, 40, 60, 80, 100}. Line styles are used to denote treatment conditions: control, pirf, pentox, and pentox & pirf. (First column): ifc = 20. (Second column): ifc = 40. (Third column): ifc = 60. (Fourth column): ifc = 80. (Fifth column): ifc = 100. Note the differences in scales between each column. Gray solid lines denote reference trajectories with constant slopes of 10×ifc/tick and 5×ifc/tick.

The final total collagen increases proportionally with ifc (Fig. 10) but not substantially across the tissue sample cases. All cases start with the same dimensionless collagen value of 1 in each patch but differ in their initial total collagen due to the percent collagen (Sect. 3.2.3). Case A starts with total collagen = 2303, case C starts with total collagen = 6627, and case D starts with total collagen = 6173. These differences in initial total collagen are imperceptible as they are three orders of magnitude smaller than the total collagen and are outweighed by the substantial amount of collagen deposited, driven by ifc.

The drug treatments are applied at the initial time, so a difference in slope in Fig. 9 compared to the control scenario indicates a treatment effect (i.e., approaching the total collagen increase rate of 5×ifc/tick compared to 10×ifc/tick without treatment).

**Fig. 10.**
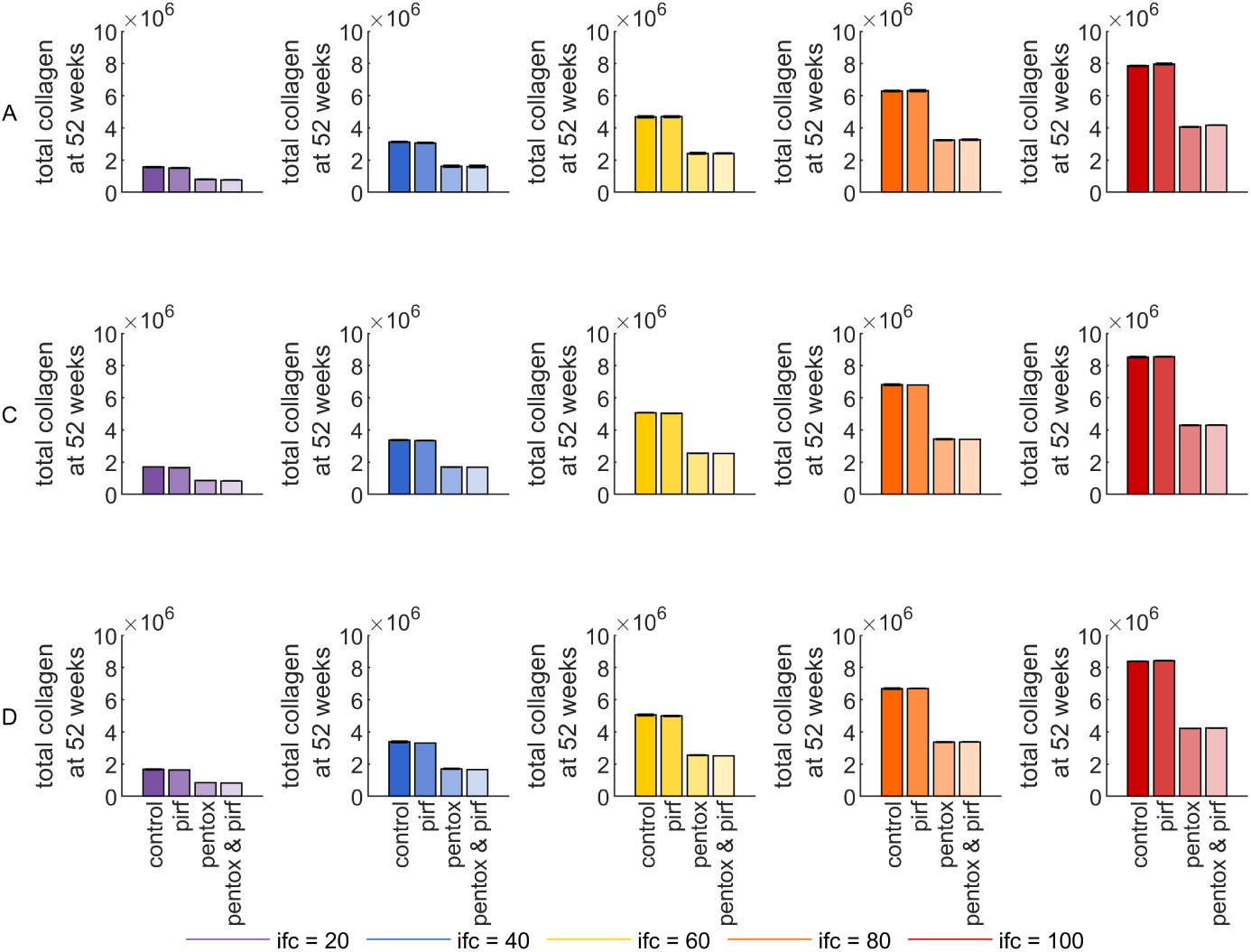
Final total collagen for 180 *in silico* experiments automated through BehaviorSpace in NetLogo for (first row) case A healthy, (second row) case C moderate fibrosis, and (third row) case D severe fibrosis lung tissue at 52 weeks of simulated time. Bars represent the mean and standard deviation of three replicates for each scenario. Colors are used to denote values of initial-fibroblast-cells = ifc = {20, 40, 60, 80, 100}. Bar color intensity is used to denote treatment conditions: control, pirf, pentox, and pentox & pirf. (First column): ifc = 20. (Second column): ifc = 40. (Third column): ifc = 60. (Fourth column): ifc = 80. (Fifth column): ifc = 100.

The slopes are similar but not identical for the control and pirf treatment scenarios. The slopes are reduced for both treatment options involving pentox. The bar graphs illustrate this more succinctly (Fig. 10), making it clear that pirf treatment in our simulations differs only to a negligible extent in the total collagen compared to the control scenario. Pentox treatment, alone or in combination, leads to a much more substantial (two-fold) decrease in total collagen. This follows directly from the fibrosis parameter pentox-myo collagen = 0.5.

#### 3.2.3 Percent collagen

The percent collagen metric (Figs. 11 and 12) monitors the extent of the migration in the domain. As cells explore the domain, they convert patches previously alveoli into collagen, representing the fibrotic invasion of the alveolar airspaces. The metric quantifies the patches that start as collagen and those where this conversion occurs.

**Fig. 11.**
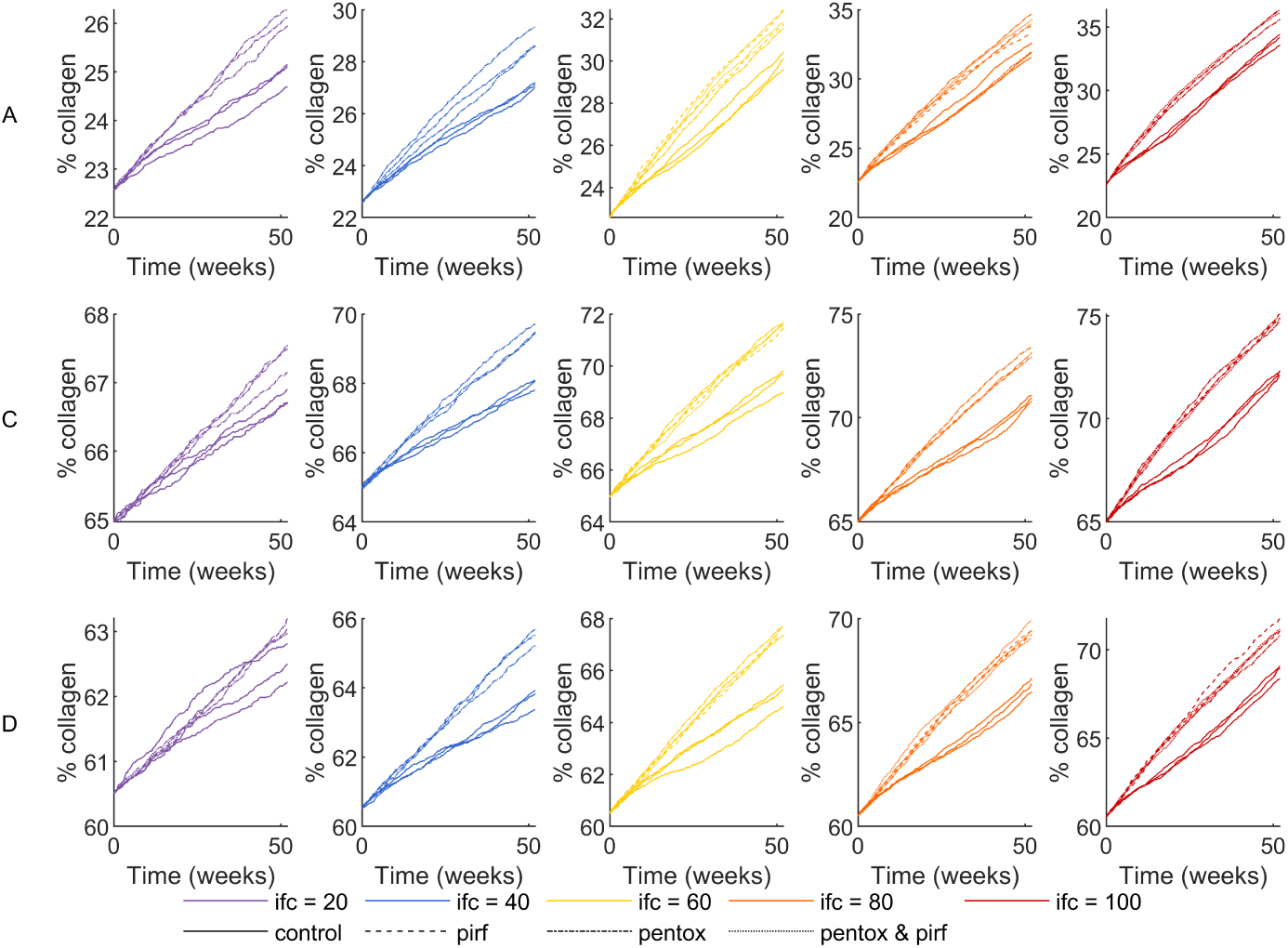
Dynamic percent collagen for 180 *in silico* experiments automated through BehaviorSpace in NetLogo for (first row) case A healthy, (second row) case C moderate fibrosis, and (third row) case D severe fibrosis lung tissue for 52 weeks of simulated time. Colors are used to denote values of initial-fibroblast-cells = ifc = {20, 40, 60, 80, 100}. Line styles are used to denote treatment conditions: control, pirf, pentox, and pentox & pirf. (First column): ifc = 20. (Second column): ifc = 40. (Third column): ifc = 60. (Fourth column): ifc = 80. (Fifth column): ifc = 100. Note the differences in scales between each panel.

**Fig. 12.**
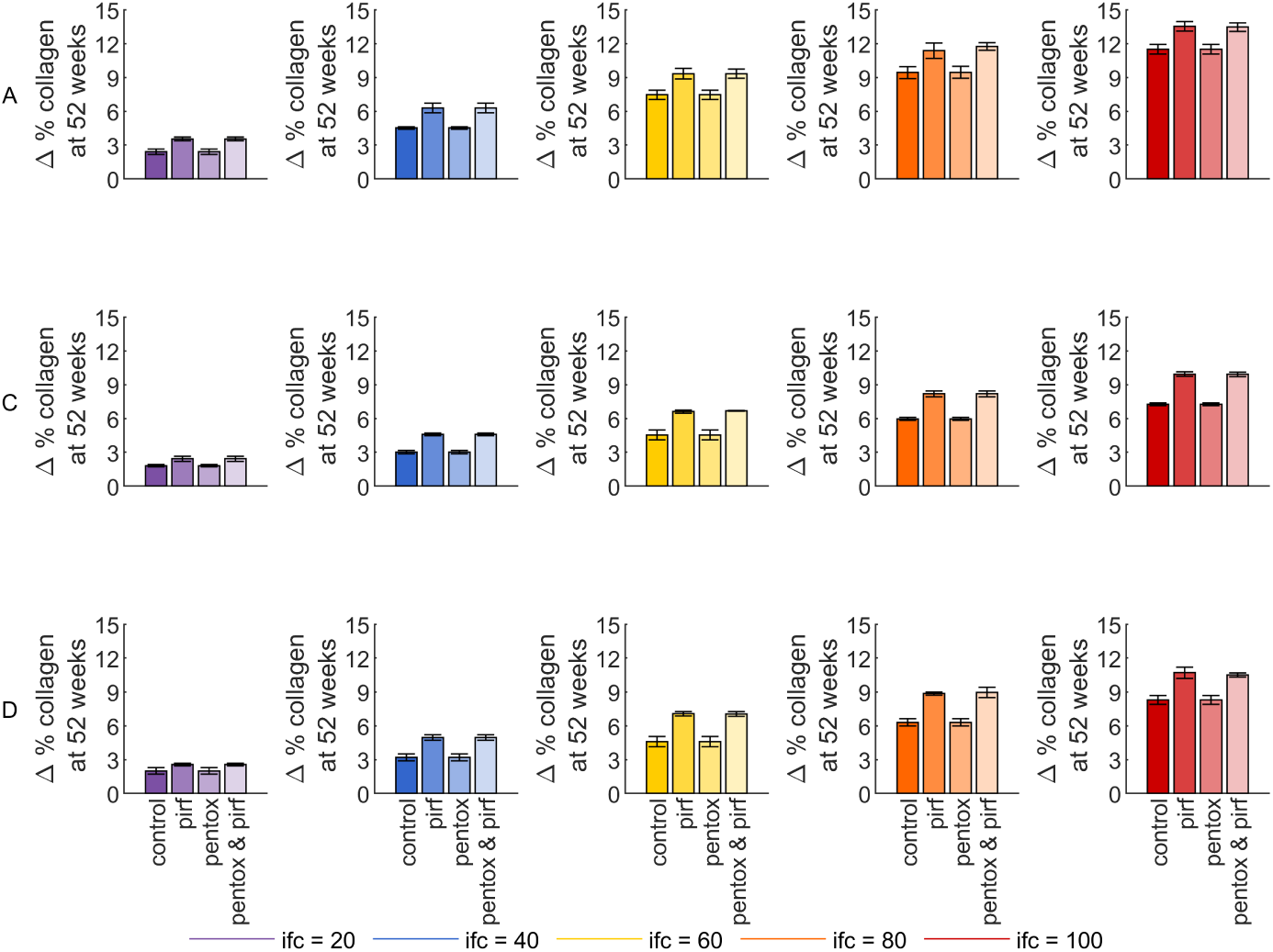
Final change in percent collagen for 180 *in silico* experiments automated through BehaviorSpace in NetLogo for (first row) case A healthy, (second row) case C moderate fibrosis, and (third row) case D severe fibrosis lung tissue at 52 weeks of simulated time. Bars represent the mean and standard deviation of three replicates for each scenario. Colors are used to denote values of initial-fibroblast-cells = ifc = {20, 40, 60, 80, 100}. Bar color intensity is used to denote treatment conditions: control, pirf, pentox, and pentox & pirf. (First column): ifc = 20. (Second column): ifc = 40. (Third column): ifc = 60. (Fourth column): ifc = 80. (Fifth column): ifc = 100.

Unlike the total collagen metric, which does not show much stochastic fluctuation over time (Fig. 9), the percent collagen fluctuates as the cells’ ability to penetrate into the alveolar region is determined probabilistically (Fig. 11). Pirf treatment, alone or in combination, has the greatest influence on the trajectories of percent collagen (Fig. 11). Ideally, it would be preferable to reduce the percent collagen (or the rate of increase) relative to the control scenario to maintain the alveolar surface for gas exchange. This behavior of the pirf treatment emerges from the combination of rules, not explicitly from any single rule in isolation. The current mechanism is to decrease TGF-β secretion, thereby reducing the chemotactic signal towards damaged inflammatory hotspots. The unintended consequence of this mechanism is the enhancement of the invasion into the alveoli patches; the reduction in the TGF-β gradient leads to the random walk mode dominating the migration of cells under pirf treatment for extended periods of time compared to control or pentox treatment, providing more opportunities for attempts to move onto alveoli. Evidence that chemotaxis is the dominant migration mode for the control and pentox treatment scenarios is visible in the contour maps for cases C and D (Figs. 5 and 6). The final spatial positions of blue and yellow levels of collagen for the control and pentox treatment scenarios correspond directly to the initial locations of TGF-β across all replicates. In contrast, the analogous maps for the pirf treatment scenarios show more diffuse collagen deposition across regions not localized to TGF-β sources, indicating that the random walk mode is the dominant migration mode for the pirf treatment.

To compare between the cases that have different starting values of percent collagen, Fig. 12 shows the final change in percent collagen for each case, ifc value, and treatment scenario. The final change in percent collagen increases with ifc and is much more pronounced for case A, where the fraction of collagen patches adjacent to alveoli is higher, giving more opportunities for successful migration into alveoli. Case A has a lower starting percent collagen, which means that the initial TGF-β injury is more concentrated (same number of sources as the other cases, but dispersed over fewer collagen patches). Thus, the response to the same magnitude of injury in fewer collagen patches is likely the major contributor to the strongly fibrotic response in case A. Pirf treatment alone or in combination yields small but noticeably higher final percent collagen values with differences more pronounced at higher ifc values (Fig. 12).

#### 3.2.4 Average collagen

The average collagen in a collagen patch is a derived quantity obtained by the ratio of total collagen to percent collagen. This value increases with ifc and decreases across the treatment scenarios in the order of control, pirf, pentox, and both (Fig. 13). The average collagen in case A is determined on a smaller number of patches containing collagen, as the final percent collagen is about half that of cases C and D (Fig. 11). Additionally, because cases C and D are more connected, cells can move more easily between adjacent contiguous regions, depositing more diffusely (i.e., in a less localized fashion) over the 52-week simulation period.

**Fig. 13.**
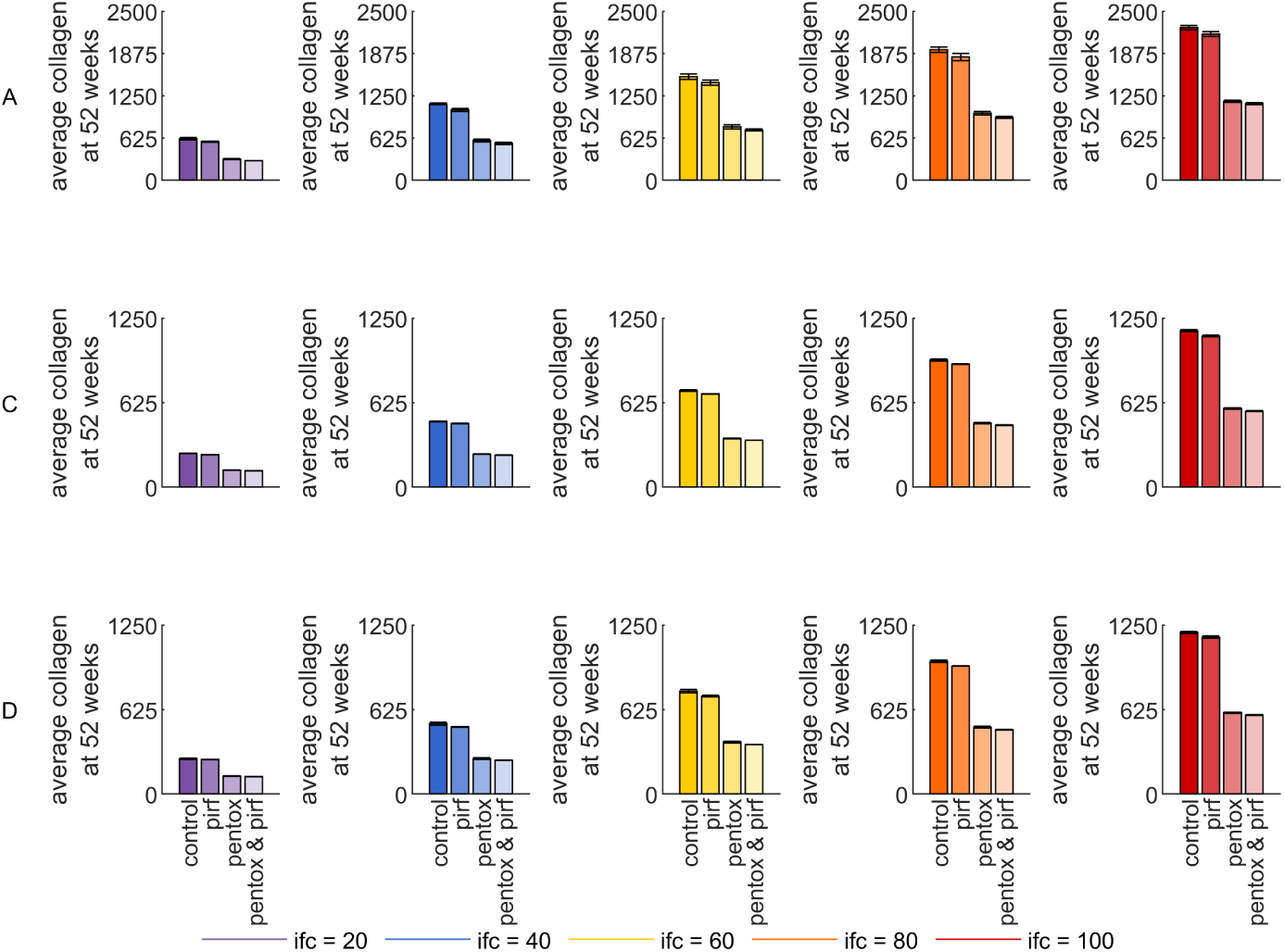
Final average collagen per patch for 180 *in silico* experiments automated through BehaviorSpace in NetLogo for (first row) case A healthy, (second row) case C moderate fibrosis, and (third row) case D severe fibrosis lung tissue at 52 weeks of simulated time. Bars represent the mean and standard deviation of three replicates for each scenario. Colors are used to denote values of initial-fibroblast-cells = ifc = {20, 40, 60, 80, 100}. Bar color intensity is used to denote treatment conditions: control, pirf, pentox, and pentox & pirf. (First column): ifc = 20. (Second column): ifc = 40. (Third column): ifc = 60. (Fourth column): ifc = 80. (Fifth column): ifc = 100. Note the differences in scales between the first row and the second and third rows.

Beyond the scalar average collagen, the contour maps of collagen show that the control scenario for Case A yields high and heterogeneous collagen spatial distribution (Fig. 4). Pirf treatment in Case A reduces the intensity and spreads deposited collagen across more patches. Pentox treatment, alone or in combination, yields much lower collagen intensity, except on isolated islands. These same trends hold for cases C and D, but at lower collagen intensities (Figs. 5 and 6).

#### 3.2.5 Max collagen

The max collagen metric (Figs. 14 and 15) is complementary to but distinct from the other collagen-related metrics. If the max collagen is close to the ratio of total collagen to percent collagen (average collagen in Fig. 13), then the domain has a more homogeneous collagen distribution. If not, then heterogeneous spikes of collagen intensity accumulate by the end of the simulation.

**Fig. 14.**
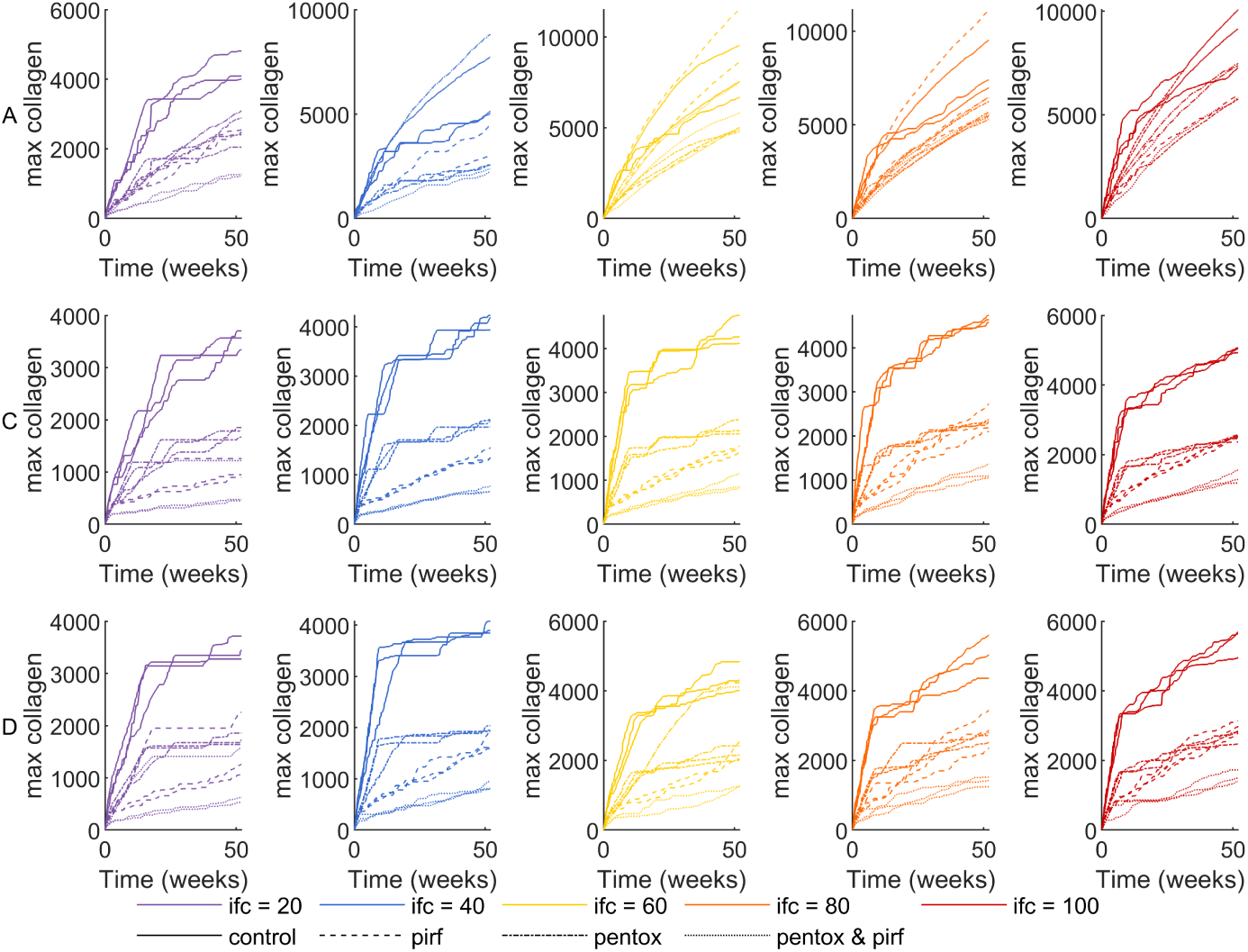
Dynamic max collagen for 180 *in silico* experiments automated through BehaviorSpace in NetLogo for (first row) case A healthy, (second row) case C moderate fibrosis, and (third row) case D severe fibrosis lung tissue for 52 weeks of simulated time. Colors are used to denote values of initial-fibroblast-cells = ifc = {20, 40, 60, 80, 100}. Line styles are used to denote treatment conditions: control, pirf, pentox, and pentox & pirf. (First column): ifc = 20. (Second column): ifc = 40. (Third column): ifc = 60. (Fourth column): ifc = 80. (Fifth column): ifc = 100. Note the differences in scales between each panel.

**Fig. 15.**
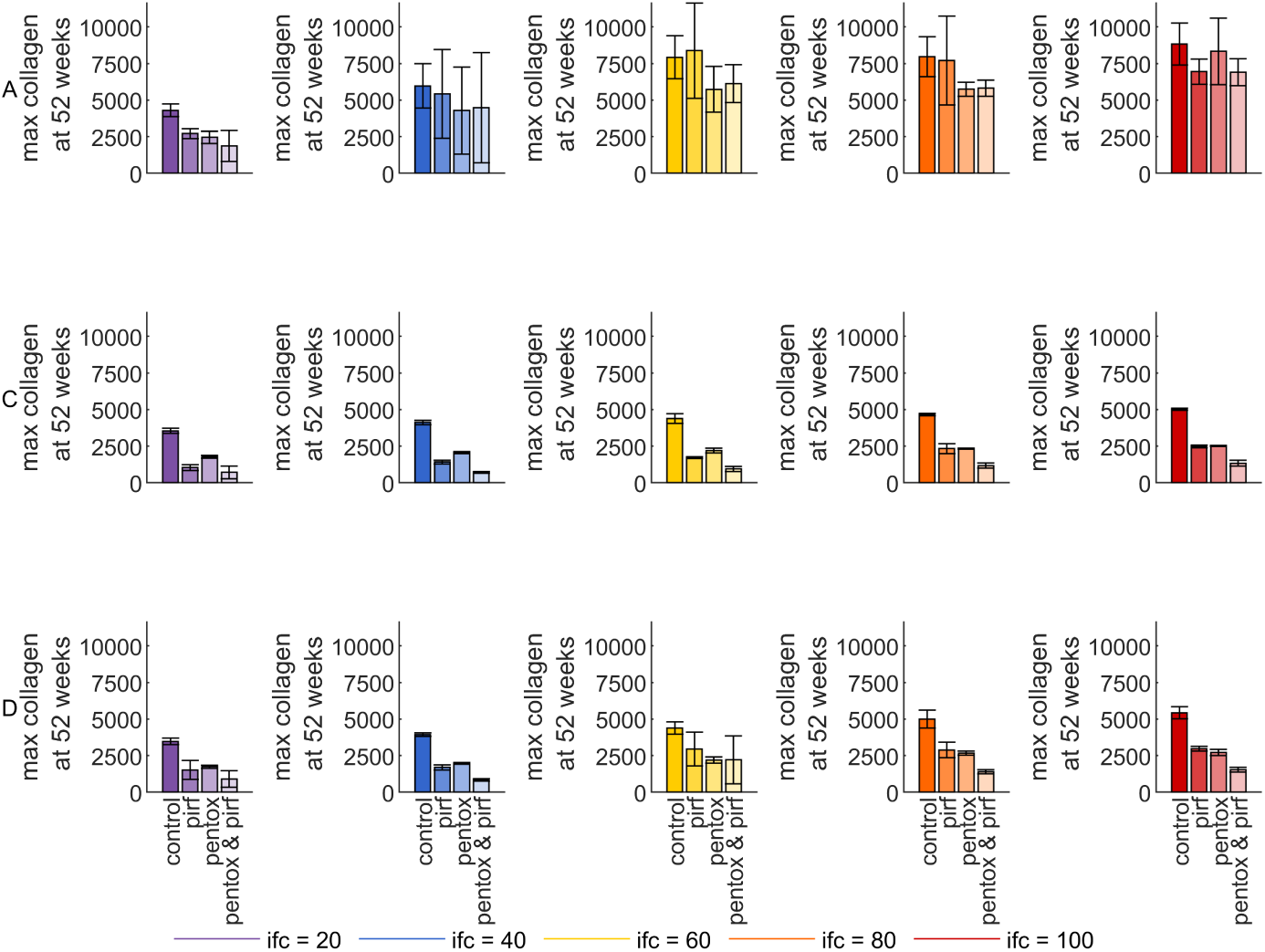
Final max collagen for 180 *in silico* experiments automated through BehaviorSpace in NetLogo for (first row) case A healthy, (second row) case C moderate fibrosis, and (third row) case D severe fibrosis lung tissue at 52 weeks of simulated time. Bars represent the mean and standard deviation of three replicates for each scenario. Colors are used to denote values of initial-fibroblast-cells = ifc = {20, 40, 60, 80, 100}. Bar color intensity is used to denote treatment conditions: control, pirf, pentox, and pentox & pirf. (First column): ifc = 20. (Second column): ifc = 40. (Third column): ifc = 60. (Fourth column): ifc = 80. (Fifth column): ifc = 100.

Max collagen depends on the treatment option (Fig. 14). The control scenario yields the highest values. Generally, the combination of pentox and pirf results in the smallest values for max collagen, particularly for the fibrotic cases C and D (Fig. 15). Notably, the max collagen for case A is impacted by the persistence of isolated fibroblast islands (Fig. 4). When those islands remain disconnected, collagen deposition by localized fibroblasts accumulates in a small region. The max collagen results for pentox alone and pirf alone are not always distinct for case A simulations (Fig. 14). For most of the simulation in cases C and D, pirf alone tends to yield smaller max collagen values over 52 weeks than pentox alone (Fig. 14). However, the effects at the final time are similar for these two treatment scenarios (Fig. 15) because the max collagen for pentox treatment tends to plateau, whereas that for pirf treatment tends to increase more consistently, sometimes surpassing the pentox plateau (Fig. 14). The plateaus for control and pentox treatment scenarios may be caused by the complete differentiation of the fibroblasts to myofibroblasts (timing for reaching plateaus seems to correspond to timing for myofibrolast dynamics in Fig. 8) in combination with having the same resulting TGF-β distribution as neither scenario affects the migration rules via TGF-β secretion.

The variation among replicates is evident in the max collagen metric, particularly for case A (Figs. 14 and 15). The max collagen results for pirf alone or in combination are generally quite variable in case D compared to the control and pentox treatment across the range of ifc values. This effect seems to be dampened at the largest two ifc values (80 and 100).

In a meta-analysis of more than 50 publications and trials involving more than 12,000 individuals with IPF [3], pirf was considered efficacious for reducing acute exacerbations compared with other treatment options. This seems consistent with the effect of a drastic reduction in max collagen at early times for simulated pirf treatment.

### 3.3 Limitations

The mechanism of pirf treatment altering TGF-β secretion did not have the expected impact on downstream signaling through indirect effects on the differentiation and fibrosis rules. The dynamics of TGF-β distributions should be considered more closely in future modeling efforts. In NetLogo, we only visualized the collagen deposition while tracking patch TGFbeta. The data for TGF-β spatial distributions were saved only at the initial and final times for each *in silico* experiment. We developed visualizations for patch TGFbeta late in our analysis and did not have the opportunity to further tune the TGF-β uptake and secretion behaviors. The current approach effectively spreads TGF-β following successful random walk moves. However, after successful chemotaxis moves, secretion occurs on the destination patch, thus further accentuating TGF-β gradients. The mechanism of pirf treatment was designed to yield a larger effective consumption rate of TGF-β (same amount of uptake and smaller amount secreted). Further model improvements could be made to reduce TGF-β gradients and localization to slow invasion by altering the target patch for secretion during chemotaxis and possibly by reducing the migration speed or the collagen secretion rate for the random walk mode when the TGF-β concentration is low. Changes in collagen secretion rate would likely lower total collagen production levels for pirf. Collagen secretion could also be made to be a function of TGF-β. Changes in migration speed may reduce invasion into the alveoli (quantified by percent collagen).

We selected ifc and drug treatment strategies applied at the initial time as the ABM input parameters to vary in our BehaviorSpace *in silico* experiments. Further experiments could vary any of the other ABM parameters listed in Table 1. Analysis of the initial TGF-β distributed at the same density, rather than the same number, across domains with different initial percent collagen may prove more realistic for the progression rate in healthy lung tissue samples.

The length and time scales were carefully chosen to be physiologically relevant. However, several ABM parameters, including those for initializations and drug treatments, were selected arbitrarily as relative quantities to compare changes over the simulation period. Calibration of these parameters to biological data and model validation are needed in the future. The collagen and TGF-β are considered in normalized units. The ABM does not account for variation or heterogeneity in the initial collagen amounts across patches, which is particularly relevant in more severe fibrosis.

Initial placement of fibroblasts and TGF-β sources did not consider the connectivity of collagen patches (leading to islands in some cases), and we did not account for the actual three-dimensional topology of the thin, curved alveolar epithelium. For simplicity in two dimensions, initial placement could be limited to collagen patches adjacent to at least one other collagen patch to prevent islands.

Although the migration speeds for fibroblasts and myofibroblasts appear as userdefined parameters, they were tested only at 1 patch/tick; adjustments would likely be needed to ensure the migration rules are robust at other migration speeds.

The ABM uses Gaussian TGF-β sources that are diminished by cell metabolism (uptake and secretion). Diffusion of TGF-β from an initial source was considered as an alternative. With a physiologically relevant diffusion coefficient of 5 × 10−7 cm2/s, the initial TGF-β was depleted within 1 hour. Simulating diffusion on time scales shorter than 1 hour and fibrosis over the time scale of years is a stiff problem.

Our ABM avoids the need for very small time steps to resolve diffusion. If Fickian diffusion is desired to be added to the model, then recurring TGF-β sources would also be needed, along with smaller time steps. The NetLogo diffuse function also would need to be constrained to prohibit diffusion into alveoli based on proximity.

More elaborate models could be developed to account for contributions from collagen degradation, influx or proliferation of new fibroblasts, cell death, and crosstalk with other cells, such as macrophages. More advanced analyses of clinically relevant fibrosis-scoring metrics or spatial statistics for point patterns could be applied to the types of *in silico* data generated in this study.

## 4 Conclusions

Through a series of *in silico* experiments, we analyzed the performance of the ABM developed in this manuscript to simulate the progression of IPF and effect of treatments. Even with the limitations of this relatively simple ABM, we determined that treatment mechanisms that directly reduce collagen secretion lowered the metrics for total, average, and maximum collagen. Treatment mechanisms to reduce TGF-β switched the migration mode from chemotaxis to random walk, lowering the maximum collagen secreted. With the current ABM settings, the role of pirf in reducing TGF-β secretion is insufficient to cascade into effects on the differentiation and fibrosis rules. Thus, the treatment scenario combining pentox and pirf is likely a better representation of the true therapeutic mechanisms of pirf, which acts on TGF-β, fibroblast differentiation, and collagen secretion.

This study also used human H&E-stained tissue sections to initialize the ABM to represent realistic tissue microenvironments. Codes for generating NetLogo domains from histological sections and bulk processing to analyze and visualize patch variables and BehaviorSpace reporter metrics were developed and shared as tools [32] for those who reuse this model or work on other NetLogo projects (see Code Availability for more details).

## Acknowledgements

This material is based upon work supported by the National Science Foundation under Grant No. DMS-1929284 while the authors were in residence at the Institute for Computational and Experimental Research in Mathematics (ICERM) in Providence, RI, during the Women in Mathematical Computational Biology program held in January 2025.

N.D.G. was supported in part by the Department of Mathematical Sciences at Rochester Institute of Technology. E.K. was supported in part by the Department of Mathematical Sciences at the University of Delaware. C.A.M. was supported in part by the University College London Department of Mathematics. A.N.F.V. was supported in part by the National Institutes of Health grant R35GM133763 and the University at Buffalo.

## Competing Interests

The authors have no conflicts of interest to declare.

## Code Availability

We have posted our codes for the agent-based model in NetLogo version 6.4.0, image processing for the human lung samples via ImageJ version 1.54, and analysis of the results and plot generation via MATLAB version R2024b in a GitHub repository [32] at https://github.com/ashleefv/LungIPFNetLogo. All *in silico* data generated is also available in the repository.

## Notes

### Competing Interest Statement

The authors have declared no competing interest.

